# SOLUBILIZER OF BACTERIAL ORIGIN SURFACTIN INCREASES THE BIOLOGICAL ACTIVITY OF C60 FULLERENE

**DOI:** 10.1101/2023.12.06.569891

**Authors:** Sergey Emelyantsev, Evgeniya Prazdnova, Vladimir Chistyakov

## Abstract

Currently, there exists conflicting data regarding the biological activity of unmodified fullerene C_60_. Various sources report its toxicity, geroprotective activity, and potential interaction with DNA. Contradictory findings regarding the toxicity of C_60_ may arise from the use of toxic solvents, as well as the influence of bioavailability and bioactivity on the preparation conditions of C60 suspensions. Furthermore, the microbiota of experimental animals can impact geroprotective activity results by releasing surfactants that facilitate substance penetration through the cell membrane.

In this study, we selected conditions for solubilizing fullerene C_60_ in a solution of surfactin, a surfactant of bacterial origin, as well as in a 2% aqueous solution of TWEEN 80, employing ultrasound. Through bioluminescent analysis using lux biosensors in E. coli MG1655, we observed that C_60_ in surfactin reduced induced genotoxic and oxidative stress. Given that surfactin enhances membrane permeability to fullerene C_60_, suspensions of fullerene in designated concentrations of surfactin can be regarded as a DNA protector and antioxidant, warranting further investigation as a promising component of novel drugs.

## 1. Introduction

Fullerenes represent the fourth allotropic modification of carbon, following diamond, graphite, and carbyne. Among the various types, the most stable and symmetrically synthesized fullerene is C_60_, commonly referred to as buckminsterfullerene. This fullerene adopts a polyhedron structure, resembling a closed sphere with 12 pentagons and 20 hexagons, with carbon atoms occupying the vertices.

In solutions, fullerenes tend to form clusters composed of multiple molecules. While C6_0_ is insoluble in polar solvents, it exhibits slight solubility in acyclic hydrocarbons. The highest solubility has been observed in aromatic solvents and their derivatives such as benzene, toluene, chlorobenzene, dichlorobenzene, as well as in carbon disulfide (1). However, the listed substances are slightly soluble in water and toxic to living cells. Thus, to prepare biocompatible aqueous suspensions of fullerenes, it is necessary to use non-toxic surfactants.

An important chemical property of the C_60_ molecule is the presence of 30 double bonds, which ensures reactivity towards free radicals. In particular, 1 molecule of _C60_ can attach up to 34 photochemically formed methyl radicals (2).

C_60_ genotoxicity studies show conflicting results and have methodological limitations due to the use of toxic solvents. According to (3), water-soluble nano-C_60_, obtained by dissolution in tetrahydrofuran (THF) followed by evaporation, is cytotoxic to cell cultures of human dermal fibroblasts, liver carcinoma and neuronal astrocytes due to ROS-mediated lipid peroxidation, without affecting DNA concentration and mitochondrial activity. However, in study (4) oral administration of a C_60_ derivative, carboxyfullerene C_60_(C(COOH_2_))_3,_ to mice from 12 months of age increased their lifespan by 11% and prevented the age-associated increase in superoxide radical generation in 24-month-old mice. mitochondria of the brain.

In (5) intraperitoneal injections of C_60_ reduced ROS formation and concentrations of antioxidant enzymes (GSH, CAT, and GPx) in rats after muscle fatigue.

Results of a study (6) on biosensors *E. coli* and *S. cerevisiae* with green fluorescent protein (GFP) – a reporter of gene expression, as well as measurement of the expression of genes associated with DNA repair in human lung cell culture A549 using RT-qPCR, micronuclear test and DNA comet method showed C_60_ genotoxicity, while the Ames test gave negative results. According to the results (7) C_60_ did not affect the frequency of mutagenesis in the cII locus of mouse pulmonary epithelial cells of the FE1-MutaTM Mouse line, the number of DNA strand breaks, but increased the level of (FPG)-sensitive sites by 1.22 times. According to (8), a mixture of fullerenes C _60_ and C _70_ does not have a mutagenic effect in the Ames test, does not induce chromosomal aberrations in Chinese hamster cell culture, and does not cause acute toxicity (dose 2000 mg/kg) in vivo in rats.

According to the results of the article (9) genotoxicity assessment using the DNA comet method showed that concentrations of 10-100 µM polyhydroxylated fullerene C _60_ (OH) _24_ have a moderate dose-dependent genotoxic effect.

The results of another study (10)of DNA damage in human lymphocytes using the DNA comet method showed that the compound C_60_[C(COOH)_2_]_3_ is not genotoxic to *Homo sapiens* lymphocytes.

In 2013, we (11) put forward a hypothesis about the ability of fullerene to be a proton carrier. The hypothesis suggested that fullerene can absorb protons and acquire a positive charge, which allows it to penetrate mitochondria. Thus, the formation of superoxide anion radical is reduced due to the gentle uncoupling of respiration and phosphorylation. This mechanism has been shown for a number of other antioxidants (12). Theoretical modeling using the DFT method showed that such a mechanism has a right to exist (13).

In 2012, an article by Baati et al. (14), which described the geroprotective properties of fullerenes. The study shows that oral administration of C_60_ dissolved in olive oil (0.8 mg/ml) in regular doses (1.7 mg/kg body weight) to rats not only does not entail chronic intoxication, but it almost doubles their lifespan effects on lifespan are mainly due to the attenuation of age-related increases in oxidative stress. Pharmacokinetic studies have shown that dissolved C_60_ is absorbed in the gastrointestinal tract and excreted within several tens of hours. The effect of unmodified C_60,_ dissolved without the use of toxic surfactants, on life expectancy emphasizes the lack of chronic toxicity.

However, in a study (15) conducted on mice, oral administration and intraperitoneal injections of C_60_ in olive oil did not significantly increase lifespan. We believe that one of the reasons for the lack of effect of C_60_ in olive oil on the life expectancy and health of mice in the study (15) may be differences in the microbiota and a lower level of production of surfactin (or other biosurfactants) compared to the intestinal microflora of rats in the experiment (14).

Surfactin (chemical formula C_53_H_93_N_7_O_13_) is a bacterial cyclic lipopeptide produced by a non-ribosomal peptide synthetase, surfactin synthetase. In particular, it is produced by the gram-positive bacteria *Bacillus subtilis* (16–18). The structure is a long chain of β-hydroxy fatty acid, the number of backbone carbon atoms ranging from 13 to 15, linked to a peptide loop of 7 amino acid residues. The most common sequence of this heptapeptide is Glu-Leu-Leu-Val-Asp-Leu-Leu, comprised of the hydrophilic amino acid residues aspartate (Asp) and glutamate (Glu), and the hydrophobic amino acids valine (Val) and leucine (Leu) (19). Surfactin exhibits antifungal, antiviral, antibacterial properties; for *E. coli* minimum inhibitory concentration is 3.125 g/l (20), is a strong surfactant (21)is able to reduce the surface tension of water from 72 to 27 mN/m at the critical micelle concentration (CMC), defined as “about 10 mg/L” by the authors (22). Surfactin has a non-specific mechanism of biological action, increasing membrane permeability due to the detergent effect exerted in concentrations from 12 to 50 μg/ml (23), which may be a consequence of the fact that the CMC of surfactin, depending on the method of its preparation, is in the range of 13-17 µg/ml (24).

In nanotechnology, surfactin has proven to be a powerful stabilizer of nanoparticle suspensions. In a study (25), it was found that silver nanoparticles, synthesized through the reduction of AgNO3 with sodium borohydride in the presence of surfactin at a concentration of 25 mg/L derived from B. subtilis (BBK006), exhibited prolonged stability for a period of 2 months. Conversely, the control experiment without surfactin witnessed rapid aggregation of the nanoparticles, resulting in the formation of large aggregates. The authors suggest that imide groups with a lone pair of electrons may play a role in the stabilization of nanoparticles, but do not exclude the stabilization of nanoparticles due to the micellar structure of the surfactin solution. Another study (26) reported the stabilization of nanoparticles through the presence of functional amino groups.

The authors (27) confirmed the possibility of using surfactin to increase the absorption of insulin in the intestine when taken orally. The use of surfactin as a surfactant replacement will reduce environmental pollution from toxic chemically synthesized solubilizers (19).

The aim of this research was to investigate the impact of C_60_ fullerene on the expression of genes linked with SOS response and antioxidant defense in an *E. coli* model, and to further examine the influence of the surfactin solubilizer on these effects.

## 2. Materials and methods

Unmodified C60 fullerene from Sigma-Aldrich, St. Louis, USA, was utilized in the experiments. Inducers of oxidative and genotoxic stress included paraquat (1,1’-dimethyl-4,4’-dipyridylium dichloride) supplied by Sigma-Aldrich, hydrogen peroxide (JSC “Ekos-1”) with special purity grade, and 1,4-dioxide of 2,3-quinoxalindimethanol (dioxidine) obtained from PJSC “Biosintez.”.

### Preparation of aqueous solutions of C_60_

Fullerene C_60_ was added to solutions of urfactin 312.5 mg/l and TWEEN 80 2 % in deionized water and sonicated with an ultrasonic dismembrator Sonics Vibra -Cell VCX 130 at a power of 30 W pulses 5/5 s for 10 min with cooling in a refrigerant to prevent boiling of solubilizer solutions and to prevent a decrease in the solubility of fullerene in them due to heating solutions above 320 K (28). The concentration of dissolved C60 fullerenes was determined by the optical density of the solution measured using a BECKMAN COULTER DU ^®^ 800 UV/ Vis spectrophotometer relative to solutions of the corresponding solvents.

### Bioluminescence test

The study of the biological properties of fullerene C_60_ was carried out on a model biological object -*Escherichia coli*.

Bioluminescent tests were performed to study antioxidant and DNA protective activities. Bioluminescent strains *of E. coli* MG 1655 carry plasmids with reporter genes *lux-CDABE,* placed under the control of promoters of oxidative and genotoxic stress and the ampicillin resistance gene *bla.* To study the biological effect of fullerene C_60_ under conditions of genotoxic stress, strains of *E. coli* MG 1655 with SOS -inducible promoters PrecA and PcolD, which detect the penetration of DNA-damaging agents into the cell, for studying oxidative stress - with promoters PkatG and PsoxS, induced by peroxides and superoxide anion radical, respectively (29). The ColD promoter has a higher induction coefficient compared to pRecA due to the lower openness of the SOS promoter *Pcda* in the absence of stress. In addition to SOS regulation by the lexA repressor, expression of the Colicin D operon is controlled by a repressor (cdr), cotranscribed with cda, but terminated earlier and preventing overproduction of colicin D in the absence of stress (30).

Dioxidine was used as an inducer of genotoxic stress at a concentration of 2.25 × 10^-5^ M, an antibacterial drug with a DNA-damaging mechanism of action due to the generation of reactive oxygen species, including superoxide (31) and, possibly, hydrogen peroxide (32); inducers of oxidative stress - hydrogen peroxide 10^-3^ M and paraquat 10^-3^ M, under aerobic conditions causing the formation of superoxide radical O^2-^ (33).

Strains were grown in liquid Luria-Bertani (LB) medium supplemented with 100 μg/ml ampicillin (34). The overnight culture, depending on the strain, was diluted with fresh LB with ampicillin to a density of 0.01-0.1 McFarland units, the culture density was measured using a DEN-1B densitometer (Biosan), then grown in a thermostat at 37 °C 1.5 h, after which 80 μl aliquots of this culture were transferred into sterile microplate cells. The control cells were cells with only solubilizers added to the culture - surfactin or TWEEN 80 (2%) (solvent control). After adding surfactin, TWEEN 80 and fullerene solutions were further incubated for 30 min in a 96-well microplate in a thermostat at a temperature of 37 ± 0.2 °C. Then inductors were added to the cells of the tablet: hydrogen peroxide and paraquat at a concentration of 10 ^-3^ M; dioxidine 2.25×10 ^-5^ M.

The microplate was placed in the FLUOstar multimodal reader Omega and incubated at 37 °С. The bioluminescence intensity was measured every 10 min.

SOS response induction factor (*I*) was calculated using the formula:

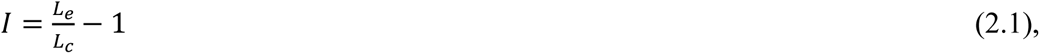

where: L_e_ – luminescence intensity of the sample with the inductor (in arbitrary units); L_c_ – luminescence intensity of the control sample (in arbitrary units).

Protective activity was calculated as

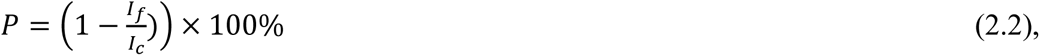

where I_f_ and I_c_ are induction factors of the surfactine solution with and without added fullerene, respectively.

Each experiment was conducted at least three times with six replicates, and statistical analysis was conducted using Student’s t-test. Confidence intervals were calculated using MICROSOFT EXCEL software (Microsoft Corporation, Redmond, WA) with a significance level of P = 0.05.

### Measuring the turbidity of bacterial cultures using a spectrophotometer

To determine the effect of a range of surfactin concentrations on the growth curves of *E. coli,* overnight cultures of bioluminescent strains were grown as for the bioluminescent test, then aliquoted in 80 μl quantities into sterile microplate wells. Solutions of surfactin and dioxidine were added in a volume of 10 µl immediately before the experiment. Measurement of the optical density of the culture at a wavelength of 600 nm was carried out using a FLUOstar device Omega every 10 min at a temperature of 37 ± 0.2 °C.

## RESULTS

### Selection of the optimal biocompatible concentration of surfactin for use as a surfactant in bioluminescent tests

To determine the biocompatible concentration of surfactin, which does not affect the growth rate of bacterial biosensors, as well as the influence of the range of surfactin concentrations in experiments with genotoxic substances, the optical density at a wavelength of 600 nm of the bioluminescent strain *E. coli* MG 1655 pRecA was measured. All surfactin concentrations used in this experiment are above the critical micelle concentration (0.017 g/L) and below the minimum inhibitory concentration, 3.125 g/L (20). In the absence of dioxidin, surfactin concentrations of 31.25 mg/l and 50 mg/l (data not shown) have no effect on the growth rate, while 100 mg/l statistically significantly reduces the growth rate by 1.5–2 times (figure 1, A). In the presence of dioxidin 2.25·10 ^-5^ M, surfactin concentrations of 31.25 mg/l and 50 mg/l had no effect on the growth rate; a 100 mg/L surfactin concentration supplemented with dioxidine reduced growth rates, possibly due to a greater increase in membrane and cell wall permeability to dioxidine (Figure 1B).

**Figure 1.**
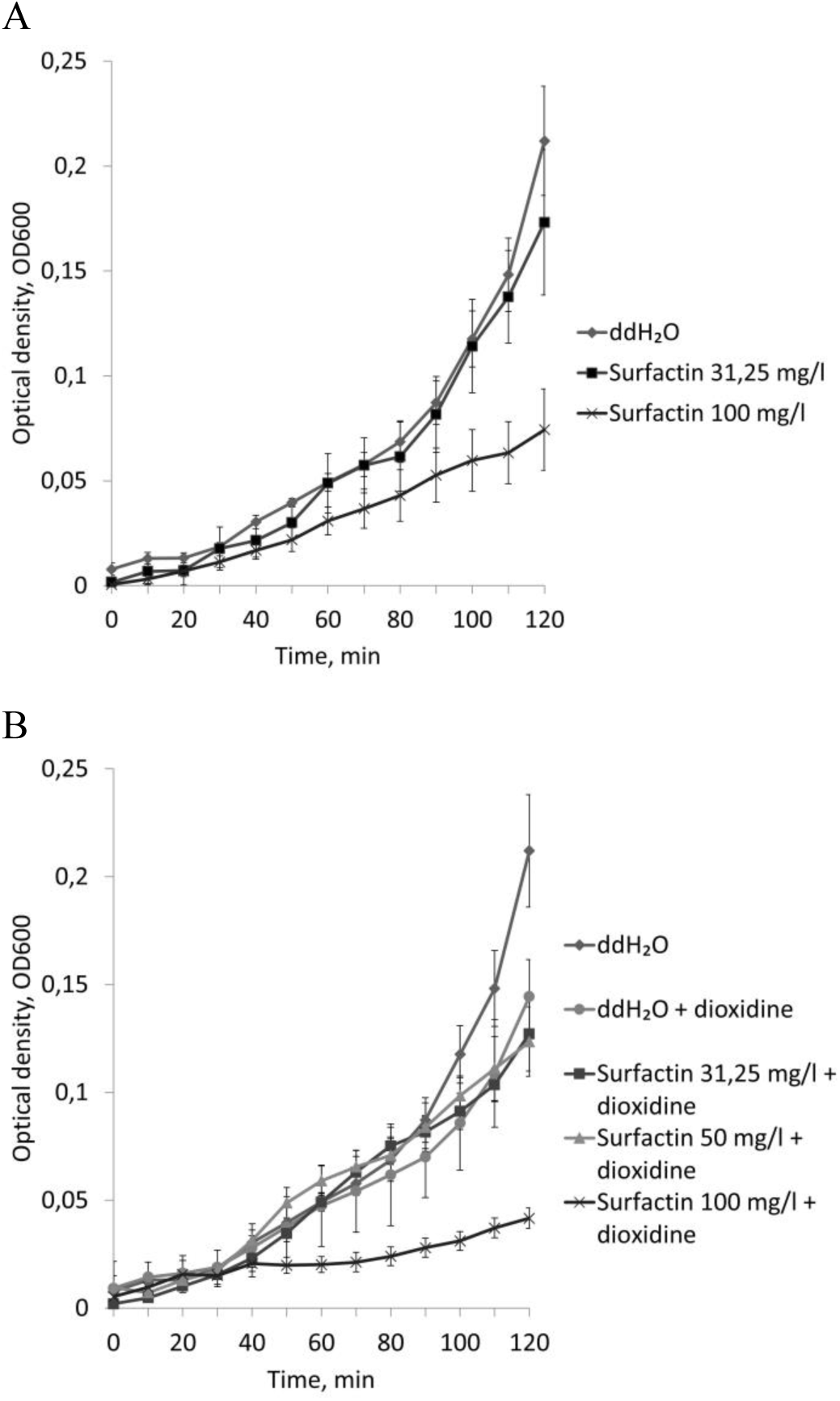
Effect of surfactin on the growth of the biosensor strain *E. coli* MG 1655 pRecA in the absence (A) and presence (B) of dioxidine.

Surfactin concentrations of 50 mg/l increased the induction of bioluminescence relative to dioxidine. Thus, increasing the concentration of surfactin within the biocompatible range does not have a protective effect, but enhances the genotoxic effect of dioxidine (Figure 2).

**Figure 2.**
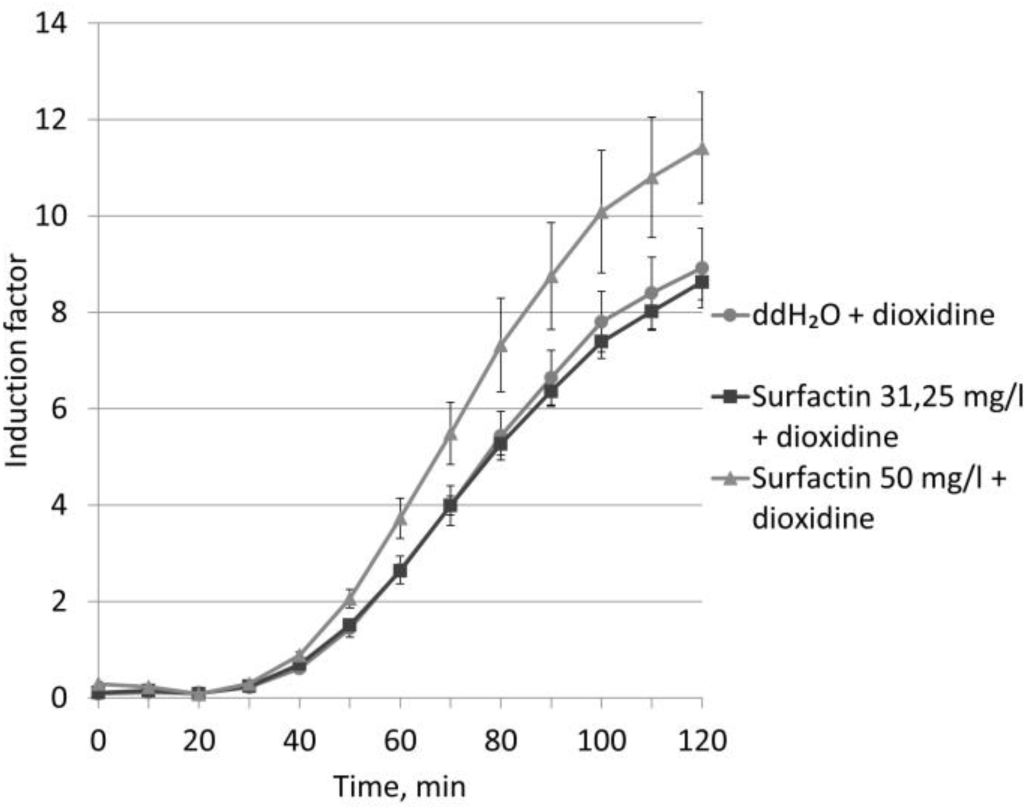
Effect of surfactin on the induction of bioluminescence by dioxidine of the biosensor *strain E. coli* MG 1655 pRecA - lux.

Based on the data obtained, for use as a solubilizer in experiments with toxic substances, it is sufficient to use a surfactin concentration of 31.25 mg/l, for the greatest increase in membrane permeability in the absence of toxicants for nanoparticles and drugs without reducing the growth rate - 50 mg/l. Surfactin reaches the micellization plateau at a concentration of 0.28 mM (290.2 mg/L) (36).

### Biological activity of solutions C _60_

For preliminary screening, C_60_ concentrations of 0.01 – 10 mg/l in TWEEN 80 (2%) and surfactin 31.25 mg/l were tested on the *E. coli* MG1655 pRecA biosensor. All tested concentrations of C_60_ in TWEEN 80 (2%) and concentrations 0.01 and 10 mg/L in surfactin had no effect on the induction factor of bioluminescence relative to the solvent control. In addition, it turned out that TWEEN 80 reduced induction factor relative to the positive control (maximum induction factor with dioxidine 20.4, while the addition of TWEEN 80 or fullerene solutions reduced the maximum induction to 12.09-13.15) (data not shown).

The surfactin concentration of 31.25 mg/l used for fullerene solubilization had no effect on the induction of bioluminescence of the biosensor strain *E. coli* MG1655 (pRecA), p <0.05 with dioxidine relative to deionized water (ddH_2_O), and subsequently the effect of C_60_ was assessed relative to this concentration (solvent control).

Solutions of C_60_ at concentrations of 1 – 10 mg/l in surfactin caused a statistically significant decrease in induction factor relative to the solvent control (Figure 3A) (induction graphs with C60 concentrations of 2.5 – 7.5·mg/l are not shown), thus demonstrating a DNA protective effect. It was found that the DNA protective effect in the concentration range from 1 to 10 mg/L in surfactin has no dose dependence (Figure 3B).

**Figure 3.**
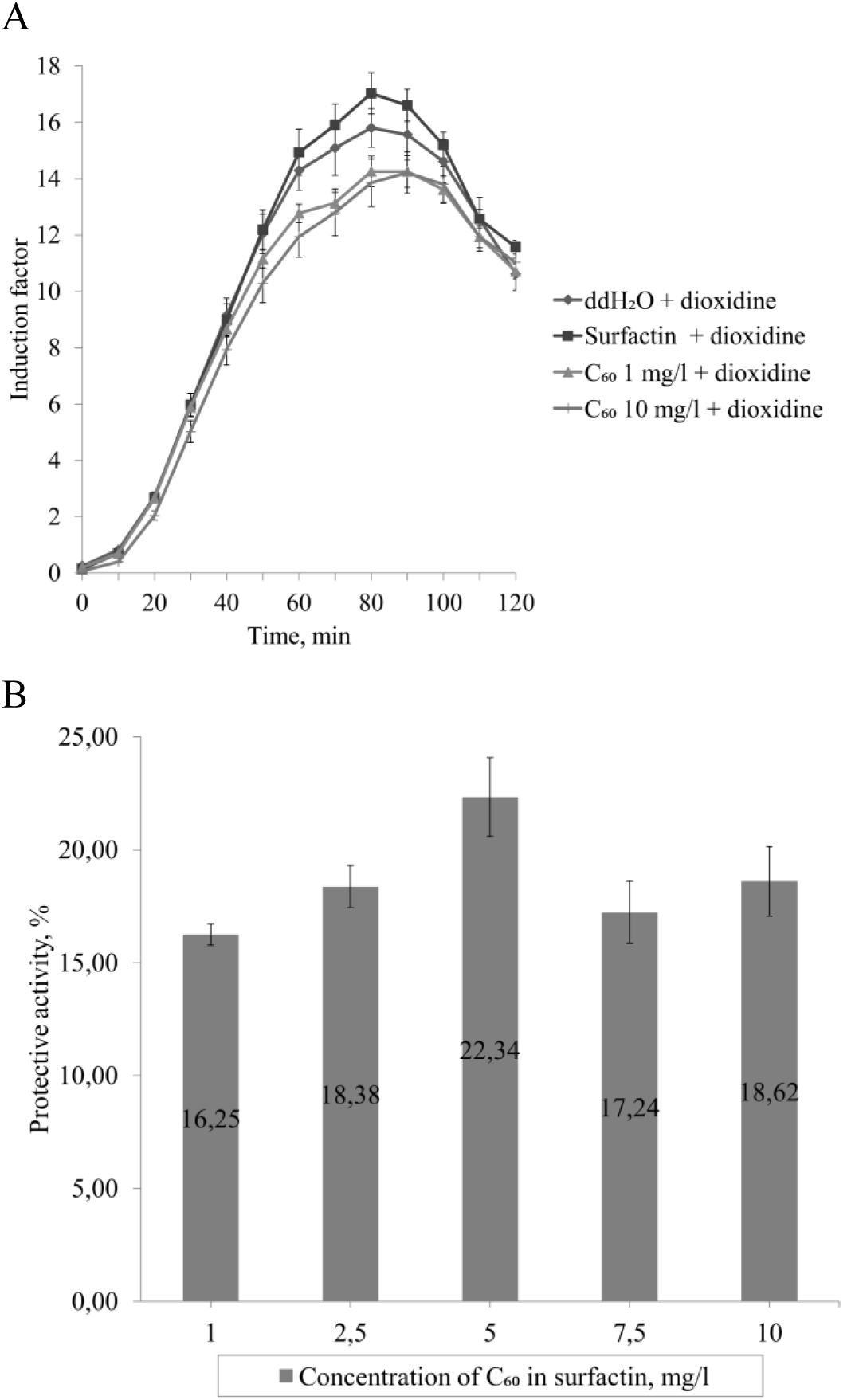
(A) Reduced induction factor caused by dioxidine 2.25×10 ^-5^ M (p < 0.05) of the biosensor strain *E. coli* MG1655 (pRecA) relative to the solvent control (surfactin) fullerene C_60_ at concentrations of 1 and 10 mg/l, solubilized in surfactin; (B) DNA protective effect of surfactin C60 concentrations during maximal induction

Thus, the biological activity of fullerene appears in surfactin solubilizer solution, but not in TWEEN 80 solution.

The results of the bioluminescent test using the *E. coli strain* MG1655 (pColD) are presented in Figure 4. Fullerene C_60_ concentrations of 0.01 and 0.1 m g/l did exhibit a statistically significant decrease in induction factor compared to the solvent control (surfactin) in the presence dioxidine, as in experiments using the biosensor *E. coli* MG1655 (pRecA) (data not shown). For the range of C _60_ concentrations 1 – 10 mg/l, a statistically significant decrease in induction factor was observed relative to the solubilizer and control (ddH_2_O) (Figure 4A) (induction factor graphs with C_60_ concentrations of 2.5·– 7.5·mg/l not shown). Concentrations of C_60_ 10^-3^ – 10^-2^ g/l in surfactin on the biosensor *E. coli* MG1655 (pColD) had a DNA protective effect close in value to that obtained on the biosensor *E. coli* MG1655 (pRecA) (Figure 4B). which also does not have a direct dose dependence. Thus, when surfactin is used as a solubilizer, C_60_ has a DNA protective effect on both biosensor strains that detect DNA damage.

**Figure 4.**
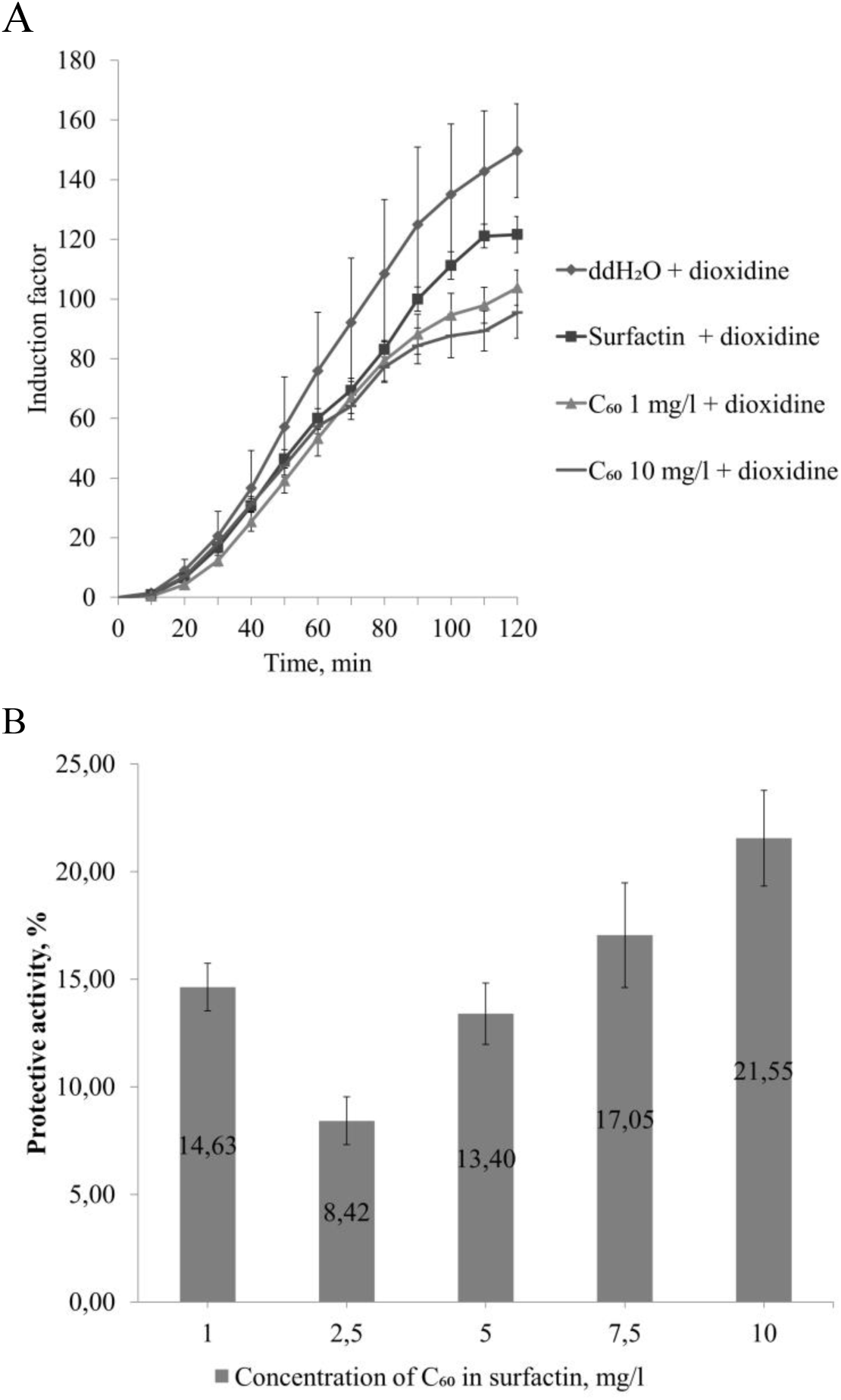
(A) Reduced induction of biosensor strain *E. coli* MG1655 (pColD) caused by dioxidin 2.25×10 ^-5^ ^M^ relative to the solvent control (surfactin) fullerene C _60_ at concentrations of 1 and 10 mg/l, solubilized in surfactin (p<0.05);. (B) DNA protective effect of C₆₀ in surfactin during maximal induction by dioxidine of biosensor *E. coli* strain MG1655 (pColD), p < 0.05.

To test the hypothesis about the antioxidant mechanism of the DNA protective effect of fullerene C_60,_ a bioluminescent test was performed using *E. coli* strains MG1655 (pKatG) and *E. coli* MG1655 (pSoxS), reacting to peroxides and superoxide anion radical, respectively. As can be seen from Figure 5A and B, concentrations of C_60_ in surfactin, which exhibit a DNA protective effect, exhibit an antioxidant effect against hydrogen peroxide 10^-3^ M. Correlation coefficient between the values of fullerene concentrations and the protective effect they provide against hydrogen peroxide during maximum induction and fullerene concentration was 0.884, which means a strong direct connection, i.e. the antioxidant effect of C_60_ is dose-dependent.

**Figure 5.**
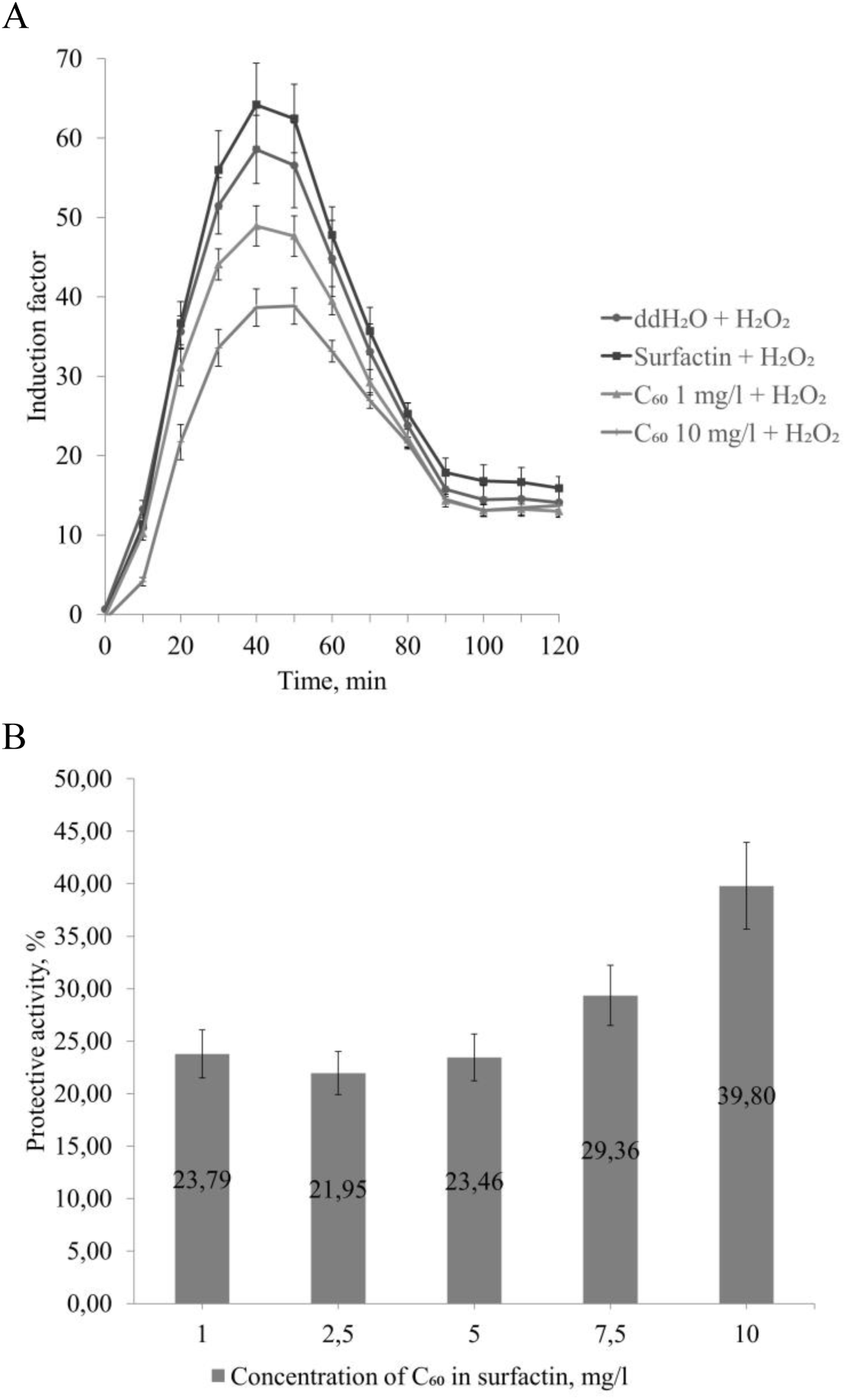
(A) Reduction of hydrogen peroxide-induced 10 ^-3^ M induction of the biosensor strain *E. coli* MG1655 (pKatG) relative to the control solvent (surfactin) with _C60_ fullerene at concentrations of 1 and 10 mg/l, solubilized in surfactin, p<0.05; (B) Protective (antioxidant) effect of _C60_ concentrations in surfactin during maximum induction, p <0.05.

Bioluminescence test with biosensor reacting to superoxide anion radical, *E. coli* MG 1655 (pSoxS) showed no effect of surfactin on induction factor with a standard inducer (paraquat (methyl viologen) 10 ^-3^ M) up to 110 min (Figure 6A). C_60_ at a concentration of 1 mg/l in surfactin did not have a statistically significant effect on the induction factor of bioluminescence relative to the solvent control and the protective effect. C_60_ at concentrations of 2.5·–10 mg/l had a decrease in induction factor relative to the control solvent (data for concentrations of 2.5·10 ^-3^ – 7.5·10 ^-3^ mg/l are not shown) and a dose-dependent protective effect (Figure 6B). C_60_ at a concentration of 10 mg/l also had a statistically significant decrease in induction relative to the control (deionized water with paraquat). The correlation coefficient between the values of fullerene concentrations and the protective effect they provide against paraquat during maximum induction and the fullerene concentration was 0.884 (as well as between the values of fullerene concentrations and the antioxidant effect they provide against hydrogen peroxide), which means a strong direct relationship, those. The antioxidant effect of C_60_ on the superoxide anion radical is dose-dependent.

**Figure 6.**
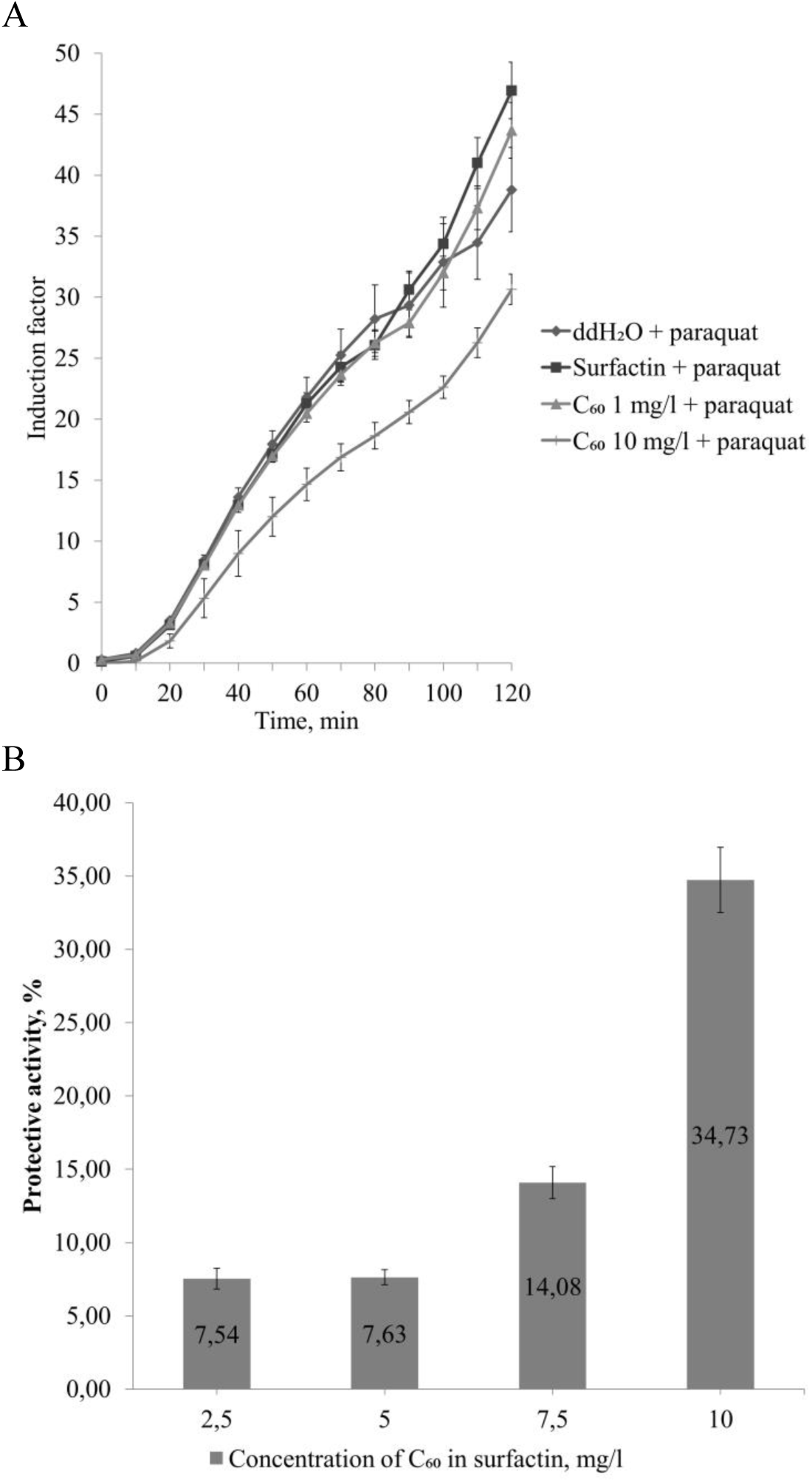
(A) Reduction of paraquat-induced induction factor of the biosensor strain E. *coli MG1655* (pSoxS) relative to the control - deionized water (ddH_2_O) and the solvent control (surfactin) with fullerene C_60_ at a concentration of 10 mg/l, solubilized in surfactin, (p<0.05); (B) Protective (antioxidant) effect of C_60_ concentrations in surfactin during maximum induction factor, p <0.05.

Thus, it can be noted that concentrations that have a DNA protective effect also demonstrate an antioxidant effect.

## DISCUSSION

From the presented data it can be seen that fullerene C_60_ has and DNA protective effect, one of the mechanisms of which is the antioxidant activity of fullerene. This is confirmed by the dose dependence of the protective effect of C_60_ on one of the ROS – hydrogen peroxide. It is also impossible to exclude the chemical interaction of fullerene with dioxidine via the Diels–Alder reaction, since the dioxidine molecule is a derivative of the heterocycle of the benzene and pyrazine rings, quinoxaline, the derivatives of which are capable of forming adducts with C_60_ (37). Perhaps the chemical interaction or adsorption of dioxidin C_60_ is the reason for the deviation from the dose dependence of the biological DNA protective effect of C_60_.

The mechanism of the protective effect against hydrogen peroxide may be the attachment of hydroxyl ion, a component of the dissociation of hydrogen peroxide (38,39); in relation to the superoxide radical, the product of partial reduction of superoxide, a hydroxyl radical, can be added to the double bonds of fullerene C_60_. It has been shown that C_60_ is capable of attaching a large number of radicals per molecule (2), including the trichloromethyl-peroxyl radical CCl_3_OO•, protecting the liver of rats from the metabolic products of carbon tetrachloride (40). It is also known that fullerenes and their derivatives, fullerenols, are also capable of acting as a powerful radical scavenger (40) and reducing ROS levels in vivo under the influence of oxidative stress inducers, such as Juglone (5-hydroxy-1,4-naphthoquinone) (41).

According to (42), the size of C_60_ aggregates and the concentration of the solution can depend on each other. In a study (43), aggregate size was found to influence biological activity. This may explain deviations in the dose dependence of the protective effect of fullerene when solutions are diluted by an order of magnitude or higher.

In our experiments, C_60_ solubilized in surfactin has a DNA protective effect, but in TWEEN 80 it does not. Therefore, it can be assumed that the bioactivity of C_60_ in surfactin is associated not so much with solubilization, but with an increase in bioavailability due to increased membrane permeability. This hypothesis explains why in the experiment of Baati et al. (14) in which fullerene C_60_ in olive oil prolonged the life of rats. Perhaps surfactin or other biosurfactants secreted by the intestinal microbiota led to an increase in membrane permeability and absorption of fullerene in the gastrointestinal tract. Surfactin-producing, in addition to potentially probiotic strains of the well-known producer *B. subtilis* (44,45) are probiotic bacteria *Bacillus clausii* (46) as well as representatives of the genus *Lactobacillus* (47) – components of the normal intestinal microflora of mammals (48) and rats in particular (49).

The lack of effect of C_60_ in olive oil on the lifespan and health of mice in the study (15) may be due to differences in the microbiota and lower levels of surfactin (or other biosurfactants) production compared to the intestinal microflora of rats in the experiment (14).

Selected for use as a solubilizer for C_60_ fullerene and to maximize the permeability of membranes for protectors, the concentration of surfactin (31.25 mg/l) is lower than the minimum concentration at which surfactin exhibits hemolytic activity (50 mg/l) (19), which opens prospects for studying suspensions of fullerene in surfactin in animal models.

Fullerene C_60_ can thus become the basis for a new type of medicine. Fullerenes can be used as antioxidants, geroprotectors, for drug delivery and in photodynamic therapy, but the above-described features of aqueous solutions, the influence of the solvent, the size of aggregates and intense lighting, especially the UV spectrum, should be taken into account.

The data obtained also indicate the potential for studying fullerene in combination with surfactin-producing bacillary probiotics in animal model organisms.

